# Heterochromatin protein 1 (HP1) of *Schistosoma mansoni*: non-canonical chromatin landscape and fitness effects

**DOI:** 10.1101/2024.03.15.585160

**Authors:** Natália S. da Trindade, Marilia Bergamini Valentini, Anne Rognon, Tiago Manuel Fernandes Mendes, Silmara Marques Allegretti, Christoph Grunau, Fernanda J. Cabral

**Affiliations:** Department of Animal Biology, Institute of Biology, University of Campinas, Campinas, Sao Paulo, Brazil; IHPE – University of Perpignan Via Domitia, CNRS, Ifremer, University of Montpellier, F-66000 Perpignan, France

**Keywords:** Heterochromatin protein 1, HP1, ChIPmentation, cercariae, sporocysts, *Schistosoma mansoni*

## Abstract

Heterochromatin protein 1 (HP1) is widespread in several organisms playing a role in control of gene expression by heterochromatin formation and maintenance of silent chromatin. *Schistosoma mansoni* is a human parasite that is responsible for Schistosomiasis, a tropical neglected disease in the tropical and subtropical areas in the world, where the intermediate host *Biomphalaria glabrata* is present. In this study we attempted to investigate if the *Sm*HP1 is enriched in *S. mansoni* chromatin in cercariae larvae stage, compared with another larvae stage sporocysts and it’s importance for *S. mansoni* life cycle progression and parasite fitness. We used ChIPmentation with commercial antibody ab109028 that passed in-house quality control. Our data show that *S. mansoni* HP1 enrichment is non-canonical with a peak at the transcription end sites of protein coding genes. We did not find strong differences in *Sm*HP1 chromatin landscapes between sporocysts and cercariae. Knock-down of *Sm*HP1 in schistosomula and *in vivo* experiments in mice unexpectedly increased parasite fitness. Our results suggest that *Sm*HP1 may influence chromatin structure in a non-canonical way in *S. mansoni* stages and may play a role in regulation of parasite fitness.

## Introduction

*Schistosomes* are trematode parasite responsible for causing schistosomiasis. It is estimated that in 2021, approximately 251.4 million people required treatment^1^. Intestinal schistosomiasis is caused by *S*.*mansoni*. The parasite has a complex life cycle that includes two hosts, snails of the genus *Biomphalaria* and humans, which are intermediate and definitive hosts, respectively^2^. During its life cycle, the parasite goes through significant stage changes, and it is known that histone post-translational modifications play important roles at each stage of the life cycle. The molecular complexity of *S. mansoni* life cycle has been revealed along years and efforts to elucidate the *S. mansoni* epigenome has given some insights about epigenetic regulation of the life cycle^3–6^ However, it is expected that coregulators will also be required for the maintenance of each stage. In several organisms, the heterochromatin protein 1 (HP1) acts as co-regulators and performs fundamental functions such as maintaining the silenced state of chromatin, DNA repair, among other diverse functions.

This protein is composed of the chromodomain (CD) and chromoshadow (CDS) domains and a linker region between them. CD recognizes and binds to methylated histone tails, while CDS is responsible for homo- and heterodimerization^7^. The region between the domains makes connections to nucleic acids and is quite variable between organisms. In contrast, the domains present well-conserved amino acid sequences among different species. HP1 is an important component essential for heterochromatin gene silencing. This function was described in model organisms such as cancer progression in *Homo sapiens, Drosophila melanogaster, Plasmodium falciparum, fission yeast* and *Arabidopsis thaliana*^8–12^. Histone modifications are associated with different chromatin states and play important roles in regulating gene expression. Particularly, it is described that in *P. falciparum* methylation of histone H3 at lysine 9 (H3K9) forms binding sites for the HP1 protein and is an important means of controlling gene expression^10,13^.

In a previous study^14^ we showed that HP1 is co-immunoprecipitated with other important DNA associated proteins such as helicases, transcription factors and methyltransferases. These results suggest an important role for HP1 in regulating gene transcription in *S. mansoni*. Geyer and colleagues also described, through *in vitro* experiments with adult worms that *Sm*CBX (Smp_179650, *Sm*HP1, *Sm* Chromobox protein homolog 5) plays a role in the parasite biology regulating oviposition^15^. We hypothesized that HP1 homolog of *S*.*mansoni* is associated with DNA and play a role in chromatin structure biology in the parasite life cycle. To test this hypothesis about the involvement of HP1 in the chromatin formation and maintenance in larvae stages, cercaria and sporocyst, were performed the ChIPmentation using an antibody against HP1 homolog used previously^14^. In addition, for investigation of the role of HP1 in parasite development, migration, fitness, and inflammatory response, we generated *Sm*HP1 *in vivo* Knock-outs. Our results suggested that *Sm*HP1 may have a function to regulate epigenetic plasticity in the parasite through the increasing of parasite’s fitness without affect host inflammatory response.

## Results

### AntiHP1 antibody ab109028 can be used for ChIP in S.mansoni

Antibody quality is an important criterion for the success and reliability of ChIP-Seq. HP1 is a conserved protein and we had shown before that antiHP1 Abcam ab109028 can be used in Western Blots with *S*.*mansoni*. A literature search for the use of ab109028 resulted in 32 publications (Suppl. Table 1) but only in one case it was used for ChIP-seq, and the efficiency of the antibody was not tested there. Thus, initial experiments were necessary to evaluate if the antibody was suitable for ChIP-Seq. We used a previously developed pipeline to test whether ab109028 (lot numbers GR38873387336-6 and -9) can be used for ChIP-Seq^16^. Essentially, the method consists of (i) performing a Western blot to assure that only the band of the expected size can be observed and (ii) doing a chromatin IP titration experiment with a constant amount of input chromatin and increasing amounts of antibody to test if saturation of available epitopes can be achieved.

We performed Western Blots on *S*.*mansoni* cercariae and sporoscysts and used *D*.*melanogaster* embryos as a positive control. Expected molecular weight of *D*.*melanogaster* HP1 is 23.194 kD (https://www.uniprot.org/uniprotkb/B6UVQ8/entry). The dimer is therefore predicted to have a molecular weight of 46 kD^14^. Such a band is observed in all samples in the Western blot (Figure 1) but supplementary bands can be distinguished. Apparently, under our relatively gentle extraction conditions and without beta-mercaptoethanol the dimers remained intact. We noticed that this behavior is not without precedent (e.g. https://www.antibodies.com/fr/hp1-alpha-antibody-a88817 ; https://www.abcam.com/en-hu/products/primary-antibodies/hp1-alpha-antibody-epr5777-heterochromatin-marker-ab109028) even for Knock-out validated antibodies (https://www.antibodies.com/fr/hp1-alpha-antibody-a88778). The results shown in Figure 1 agree also with previous western blotting^14^ which had demonstrated that *Sm*HP1 can form dimers in solution, showing a band of approximately 56 kD.

**Figure 1:**
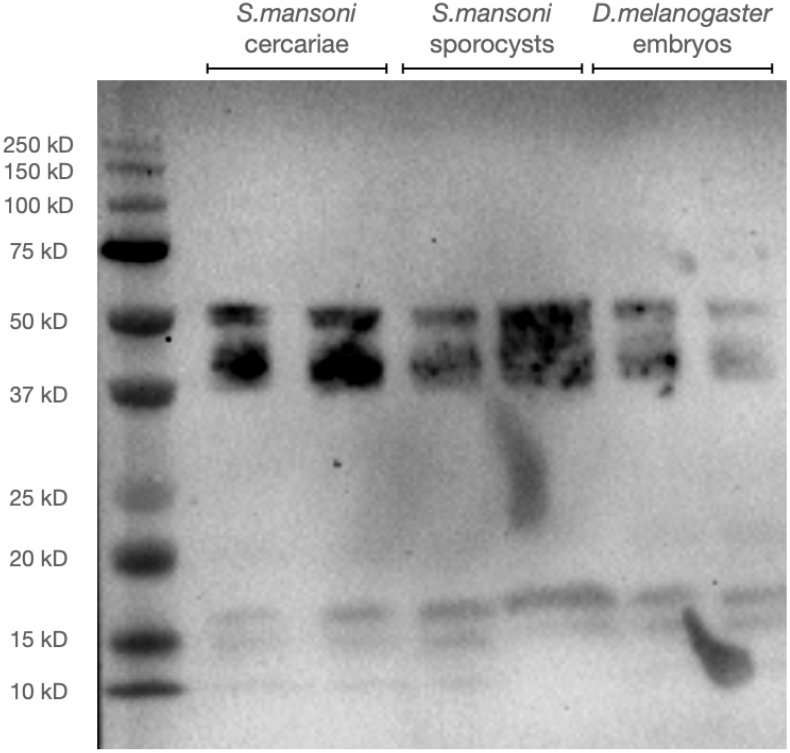
Western blot on crude protein extracts of *S*.*mansoni* cercariae and sporocysts (left and middle), and *D*.*melanogaster* embryos (right). All lanes in duplicates. Left lane molecular weight marker.

We then proceeded to ChIPmentation titration of the antibody using chromatin from *S*.*mansoni* cercariae. ChIPmentation is a streamlined method of crosslink Chromatin Immunoprecipitation (ChIP) that uses Tn5 for integration of adaptor for library amplification (Tagmentation). A constant amount of chromatin corresponding to 160,000 cells was incubated with 0, 2, 4, 8 and 16μl of ab109028 during the ChIPmentation procedure and input recovery was measured using qPCR on 2 arbitrarily chosen loci: *Sm*-alpha-tubuline and *Sm*-28S-rDNA. Results are shown in figure 2, indicating that above 8 μl antibody saturation is achieved.

**Figure 2:**
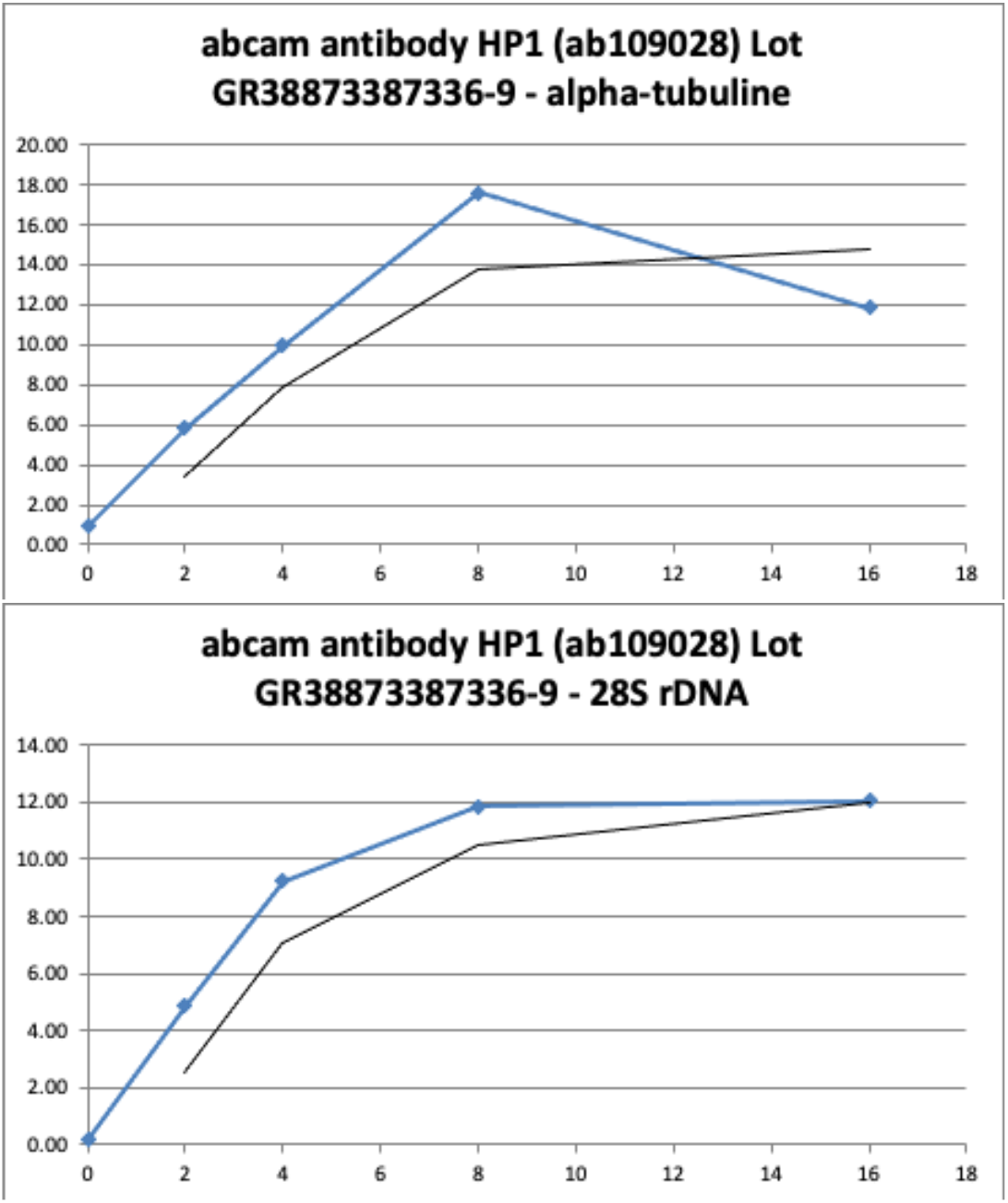
Results of titration of antiHP1 antibody. X-axis indicates amount of antibody in μl, Y-axis input recovery in %. Upper panel for Sm-alpha-tubuline, lower panel for Sm-28S-rDNA. Saturation is achieved with 8 μl of antibody.

Based on this initial testing we decided to proceed to ChIPmentation with 8μl of antibody.

### ChIP-Seq metagene profiles in *S*.*mansoni* peak around transcription end sites (TES)

After having firmly established that ab109028 was suitable for ChIP we proceeded to ChIPmentation on cercariae and sporocycts of *S*.*mansoni*. We hypothesized that HP1 could establish a heterochromatic structure in the cercariae that are transcriptionally inactive. To investigate the distribution of HP1 with relation to known genomic features we produced metagene profiles around protein coding genes. In contrast to what was observed in other species, *Sm*HP1 shows enrichment around the TES (figure 3) in both, cercariae and sporocysts.

**Figure 3:**
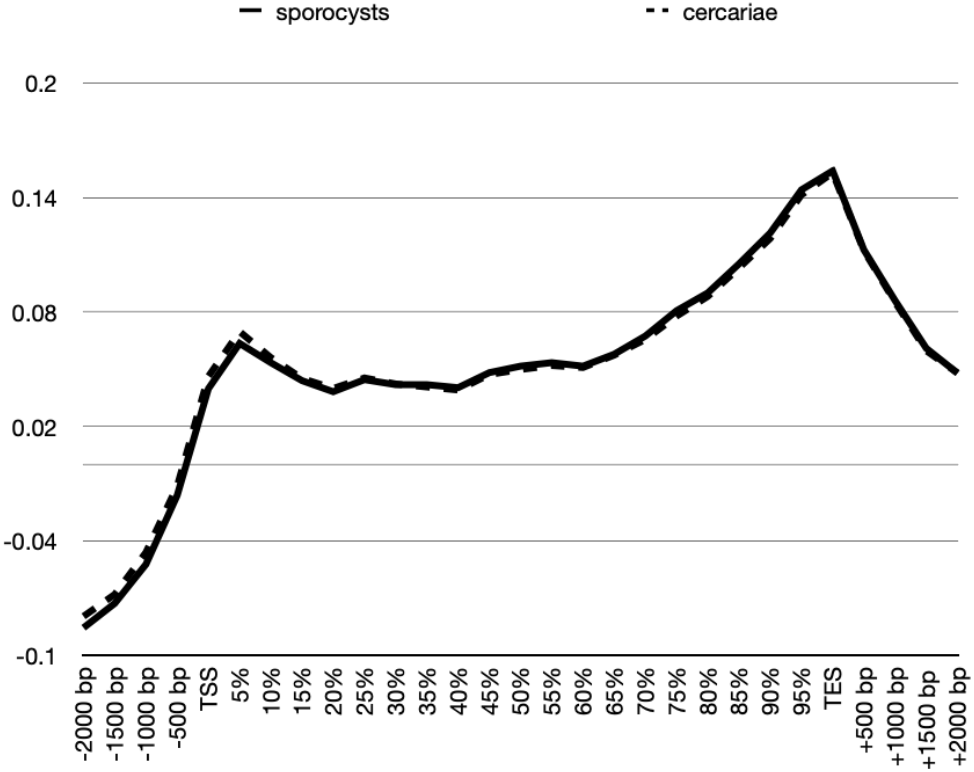
Metagene profiles of *S*.*mansoni* sporocysts (solid line) and cercariae (dotted line). X-axis : relative position around genes, Y-axis : average log(RPKM) of two replicates, TSS: Transcription start site, TES: transcription end site

This unexpected result and the fact that the current study is the first analysis of HP1 distribution and there are therefore no precedents to compare with, prompted us to test the ChIPmentation procedure with antiHP1 ab109028 on the well characterized model species *D. melanogaster*.

### AntiHP1 antibody ab109028 delivers canonical ChIP-Seq results in D. melanogaster

It might be argued that the observed metagene profiles in *S*.*mansoni* are due to a peculiar nature of the antibody that remained unnoticed. We therefore performed ChIPmentation experiments with *D*.*melanogaster* embryos, a species for which data of ChIP-Seq experiments are available. We used the query terms “hp1 chip drosophila” to search the NCBI SRA and obtained 186 results belonging to 55 BioSamples. We downloaded ChIP-Seq data for WT embryo replicate 1 and 2 (NCBI SRA: SRS6795886, SRS6795887) corresponding to ENA SRX8497063 and SRX8497065 and data for input (NCBI SRA SRS6795884, ENA SRX8497062). The NCBI SRA entry states that 2 μg antiHP1, Developmental Studies Hybridoma Bank C1A9 had been used. ^17^

We also downloaded ChIP-Seq data for fly heads that had been produced with the same antibody^18^ (NCBI BioProject PRJNA490276): ENA SRR7817540, SRR7817541, SRR7817542 for 3 ChIP-Seq replicates and ENA SRR7817573 for the input.

We then performed ChIPmentation with antibody ab109028 on fruit fly embryos under the same conditions as for our *S*.*mansoni* samples. After sequencing, we processed the published data and our experimental data as described for the *S*.*mansoni* samples. For SRA data of embryos, ChromstaR ^19^did not manage to construct a differential model probably due to relatively low enrichment of the reads. We resorted therefore to read counts distributions around metagenes (log(RPKM) instead of log(obs/exp), shown in figure 4). While it is interesting to note that there is a small decrease in adult flies compared to embryos at the TES (figure 5), we did not observe strong differences between the profiles generated based on the data of the two independent earlier studies and our experiment.

**Figure 4:**
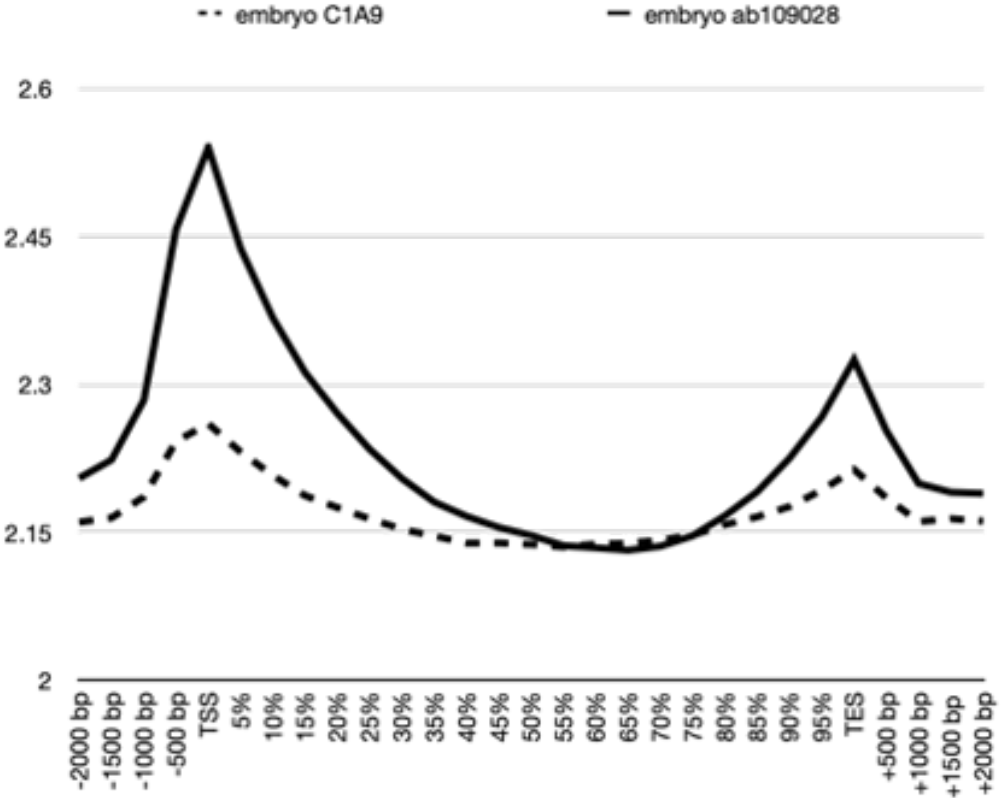
Metagene profiles of *D*.*melanogaster* embryos (solid line) produced in this study with antibody ab109028, and previously published data on embryos (dotted line) with antibody C1A9. X-axis : relative position around genes, Y-axis : average log(RPKM) of two replicates (the values can therefore not be compared directly to y-axis of figures 3 and 4)

**Figure 5:**
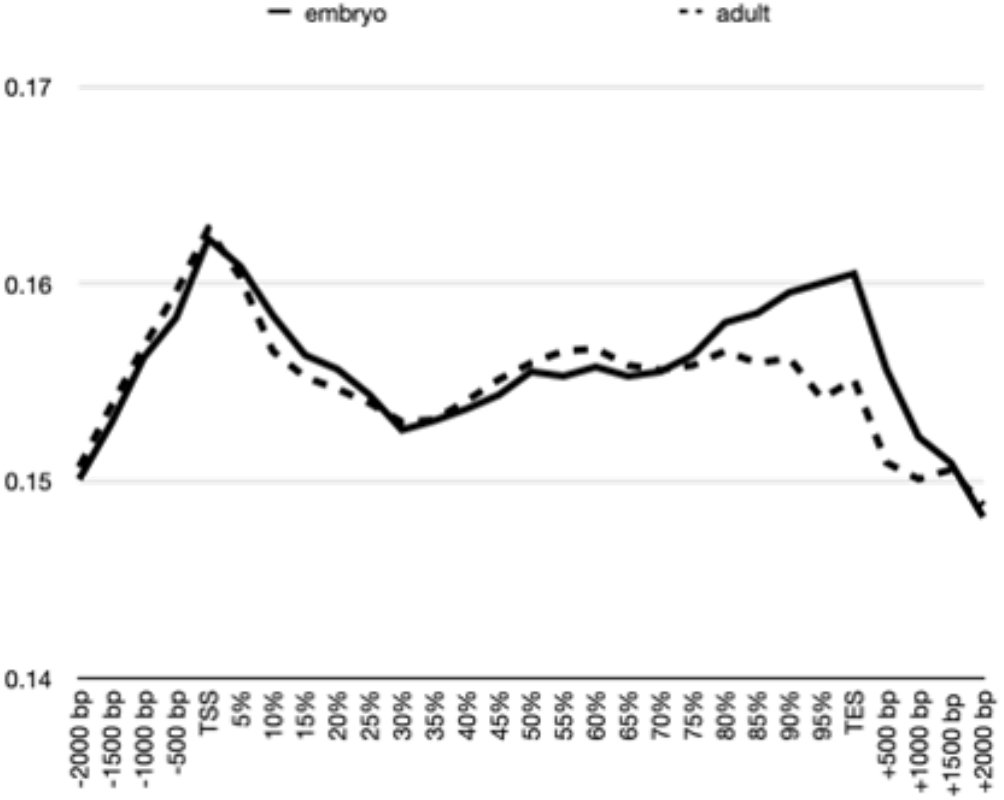
Metagene profiles of *D*.*melanogaster* embryos (solid line) produced in this study with antibody ab109028, and previously published data from another laboratory on adult flies (dotted line) with antibody C1A9. X-axis: relative position around genes, Y-axis log(observed/expected) of two replicates

Therefore, we conclude that antibody ab109028 delivers canonical ChIP-seq profiles around genes with the fruit fly as a model. Consequently, we have no reason to believe that the non-canonical HP1 chromatin landscape around genes in *S*.*mansoni* is due to the antibody quality or the ChIPmentation procedure.

### There are few regions with differential enrichment of HP1 between cercariae and sporocysts

We used ChromstaR to identify 154 regions with differences in HP1 enrichment between cercariae and sporocysts (Suppl. File 1) . However, these regions are small and visual inspection of HP1 landscape suggested that HP1 enrichment occurs over broader regions. We therefore applied MACS bdgbroadcall on combined uniquely aligned ChIP-Seq reads, independently for sporocysts and cercaria. Results are in table 1.

**Table 1:**
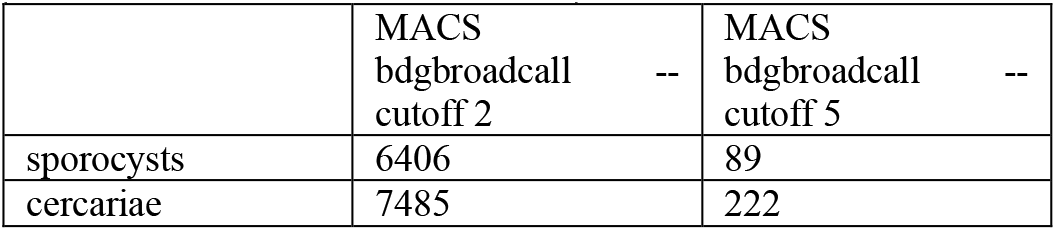
Number of broad peaks in cercariae and sporocyst larvae using different pvalue scores from MACS2, score 5 means pvalue 1e-5.

We then used DiffBind for the identification of differential enrichment between sporocysts and cercariae. Based on the conservative peakcall at score 5 we identified 36 differentially enriched regions that separated sporocysts and cercariae into 2 clusters (figure 6).

**Figure 6:**
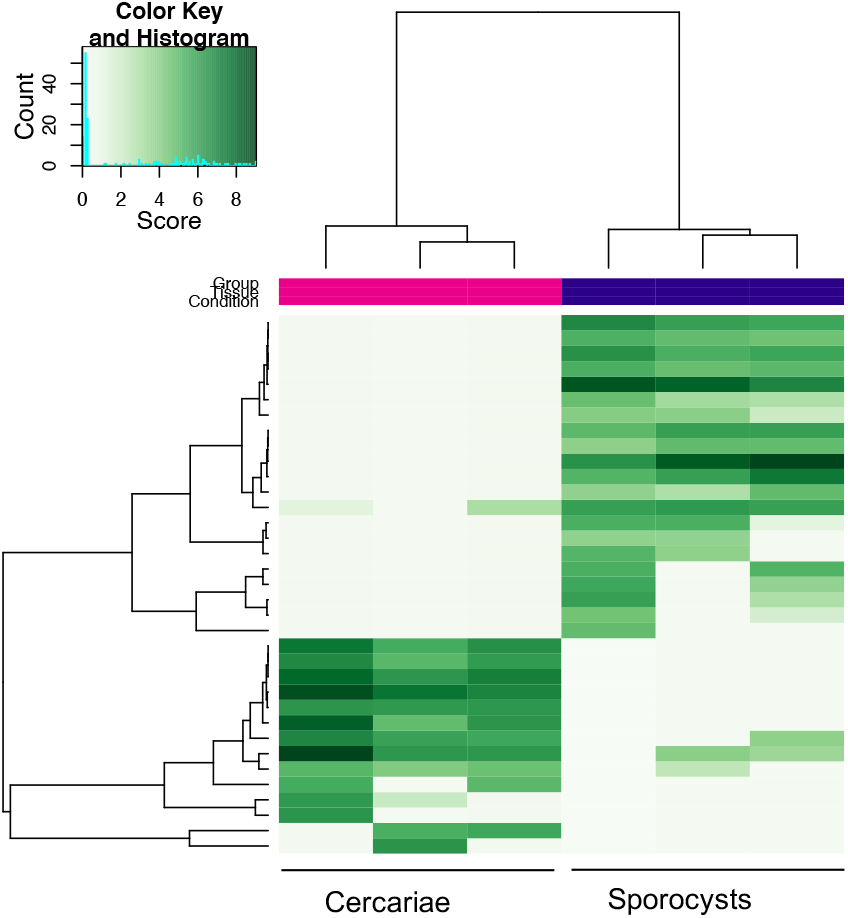
Clustering of Cercariae and sporocyst larvae based on differential HP1 chromatin enrichment

Since this number is relatively low, we wondered whether the number of identified differences could have been obtained by chance alone. We used bootstrapping through random segments that matched the 222 broad cercariae HP1 peaks in number and total span and used DiffBind for identification of differential enrichment. Bootstrapping through these analyses found in average 61.5±6.8 differential peaks. Consequently, we conclude that the differences identified by DiffBind identified on broad regions lack the statistical power to be confident about their occurrence other than by accident.

We then pursued with visual inspection on IGV, to search for regions with HP1 enrichment. Results are shown in figure 7. We observed peaks of HP1 spanning the gene Smp_330520 in sporocysts and cercariae with higher enrichment of HP1 in Sporocysts. This region overlaps also a differential peak identified by ChromstaR (figure 7a). We also identified a large region of roughly 22 kb spaning the gene Smp_344610 in which HP1 is enriched in cercariae (figure 7b). It is noteworthy that this enrichment is not observed in the input (genomic DNA) excluding therefore genetic differences between the samples in these regions. Expression in https://v7test.schisto.xyz/smansoni-v7/ revealed mRNA presence of Smp_330520 in cercariae but not in sporocysts, and very low expression of Smp_344610 in both stages.

**Figure 7:**
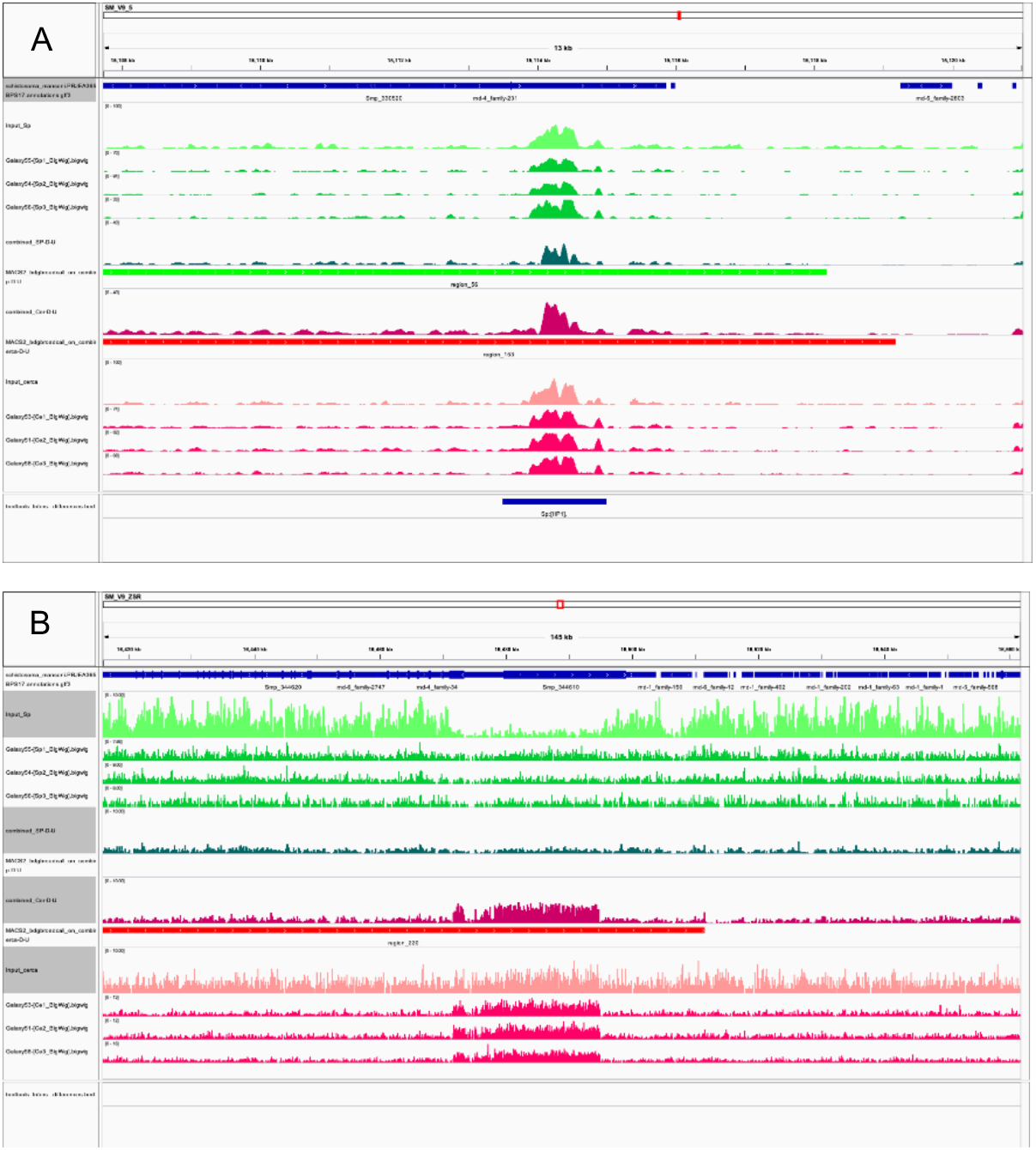
Genomic regions identified by visual inspection with differential enrichment of SmHP1 in cercariae and sporocyst larvae. IGV Screeenshots of two genomic fragments that contain diferentially enriched regions : A. SM_V9_5 15,108-15,122 kb spanning Smp_330520; B. SM_V9_ZSR 16,420 – 16,560 kb spanning Smp_344610. Lanes from top : GFF annotation (blue), Input sporocysts (light green), HP1 enrichment in 3 replicates for sporocysts (green), enrichment of combined sporocysts replicates (dark green), MACS broad peak call sporocysts (green), enrichment of combined cercariae replicates (dark magenta), MACS broad peak call cercariae (red), Input cercariae (pink), HP1 enrichment in 3 replicates for cercariae (magenta), ChromstaR detected differential enrichment (blue)

In summary, our finding cannot formally exclude that there are differences between sporocyst and cercariae in HP1 enrichment, but differences are small.

### In vivo knock-down of SmHP1 increases parasite’s fitness in mice

Given the unexpected distribution of *Sm*HP1 over the genome and absence of important differences between two developmental stages we wondered if *Sm*HP1 played any role in the parasite *in vivo*. We designed a experiment to perform the knockdown of *Sm*HP1 by RNA interference in schistosomula. Decrease of expression was small (Suppl. Figure 1) compared with control dsmCherry and wild type schistosomula. The *in vivo* results are shown in the Figure 8. Parasite burden showed that parasite was able to migrate, develop and to achieve the correct destination that is portal hepatic and mesenterial veins, despite dsRNA employed and schistosomula injection. To evaluate infection success and parasite fitness, we measured granuloma and egg count/ feces gram using Kato-Katz method^20^. The number of granulomas and area were the same for all tested conditions, suggesting that knock-down of *Sm*HP1 does not affect the granulomatous response from host. Surprisingly, the parasite number of eggs/feces gram were roughly 2 times higher in the knock-down.

**Figure 8:**
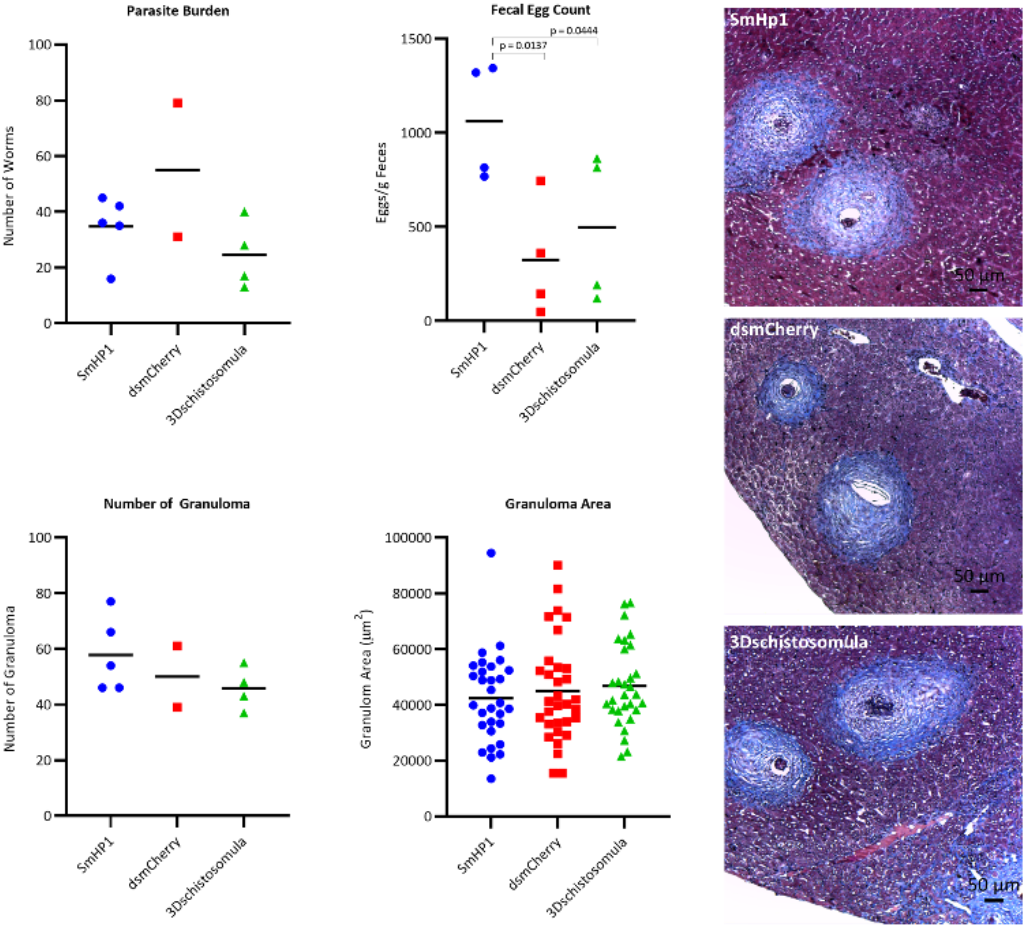
Functional *in vivo* studies in mice injected with (i) dsRNA incubated HP1 3 days schistosomula (blue) compared to (ii) mock treated dsRNA gene (dsmCherry), and (iii) wild type. Parasite burden, fecal egg count, granuloma number and area were measured. Significant differences were only found for fecal egg count which increased for the knock-down parasites.

## Discussion

Recent years have seen an important increase in our understanding of chromatin structure of *S*.*mansoni*. However, this knowledge stems almost exclusively from work on histone modifications and DNA methylation. To widen our view to other chromatin proteins we focused here on the HP1 homologue.

Our results of antibody titration show you that antibody ab109028 is suitable for ChIP-seq experiments on *S*.*mansoni*. Our report shows for the first time the HP1 profile for this species.

There are several works that have focused on transcription and translation in cercariae, absence of transcription but use of mRNA from previous stages, low rates of translational until infection and schistosomula migration when transcription is activated. ^21–23^Since cercariae is transcriptionally silent and poised for transcription ^4^, we reasoned that they might have a tighter heterochromatin formation and differential HP1 occupancy, but our results suggest the opposite: HP1 presence (and presumably heterochromatinization) might not be advantageous if it is necessary to rapidly activate transcription after infection.

HP1 is a conserved member of the large family of chromatin proteins, and it is believed to fulfil conserved functions in different organisms. For example, in Drosophila there are 5 isoforms of HP1 and it is mainly located in heterochromatic regions rich in H3K9me3^17^. however, in Drosophila HP1 is found preferentially associated with heterochromatin, polytene chromosomes and the X chromosome. The study also suggested that the formation of heterochromatin occurs in parallel with the transcriptional activation of the nuclei and expression of different genes^24^. Previous work using RNA interference technology have found that HP1 silencing was directly related to chromosomal defects^25. 24^The distribution of HP1 throughout the *S. mansoni* genome does not follow a pattern observed for other organisms suggesting that its function is also different.

Our in vivo findings indicate an increase of fitness through an increase of oviposition. Even with a moderate decrease of *Sm*HP1 expression it was possible to observe an increase of oviposition. This goes in line with earlier results were alteration of oviposition was shown in *in vitro* adult worms’ culture with siRNA against *Sm*CBX/*Sm*HP1 (Smp_179650). In this work, it was also demonstrated that *Sm*CBX/*Sm*HP1 interacts with *Sm*mbd2/3 colocalizing together with schistosome neoblasts and reproductive tissues, suggesting a role on reproductive organs of parasite^15^

Our results are another step towards the better understanding of the role of *Sm*HP1 in chromatin structure, gene expression and parasite fitness. It shows that results from model organisms, albeit tremendously useful in many cases, cannot be simply extrapolated to any other organism. *Sm*HP1 seems to play according to rules that are yet to be discovered.

### Materials And Methods

#### Extraction of sporocysts and cercariae

Sporocysts were obtained from *Biomphalaria glabrata* strain *Bg*GUA, 6 months infected with *S*.*mansoni* strain *Sm*DFO. Dissection was performed at room temperature. The snails were placed individually in large petri dishes and water was added. With the help of smaller glass plates, the snails were crushed, and fragments and tissues were removed. The sporocysts were transferred to new petri dishes and the membrane surrounding them was removed. Then, the hepatic pancreas was removed, and two samples were obtained for each snail, sporocysts free of *Biomphalaria* tissues and sporocysts that may be contaminated with *Biomphalaria* tissues. Sporocyst samples were stored at -20°C

Cercariae strain *Sm*VEN were obtained from snails *B. glabrata*. The snails were placed in pots of water and exposed to artificial light for 2 hours. Next, the Cercariae were counted and divided into aliquots with 1000 Cercariae each. They were then stored at -20°C.

Experiments were done according to the national ethics standards. The IHPE animal facility holds agreement number F6613601 and has authorization APAFIS #39910-2022121915564694 v2 for the routine production and shared used of *S*.*mansoni* larvae.

#### ChIPmentation *S*.*mansoni* cercariae, sporocysts and drosophila

ChIPmentation Kit for Histones (Diagenode, Cat. No. C01011009) was used. Approximately 1000 cercariae, 1 sporocyst and a corresponding amount of drosophila embryos were used for each ChIPmentation reaction. Samples were thawed, resuspended in 500 μL of HBSS, crushed for 1 minute with a sterile pestle, and then left at room temperature for 3 minutes. After this, 13.5 μL of 37% formaldehyde was added and gently homogenized for 10 minutes at room temperature. To stop the cross-link reaction, 57 μL of glycine from the Diagenode kit was added and left under agitation for 5 minutes. We then proceeded according to the suppliers manual. Chromatin was disrupted by sonication using the Bioruptor Pico with 5 cycles of 30s ON and 30s OFF. ChIPmentation were done on an IP-Star pipetting robot according to the pre-established protocol with the modification of washing time to 20 minutes. Antibody titration was done as described in^16^ with 0, 2, 4, 8 and 16μl antibody to obtain the saturating quantity and finally 8 μL of antibody HP1 AbCam (ab109028, lot GR38873387336-9) were used for each reaction. Input libraries were generated as described earlier and optimal number of library amplification cycles was determined by qPCR as described in the same protocol^26^.

Primers and low molecular weight fragments were removed from the libraries with AMPure beads on the IP-Star and the quality and quantity of the sequencing libraries were checked with a BioAnalyzer High Sensitivity DNA Assay. Sequencing was done by the BioEnvironnement core facility on an Illumina NextSeq 550 as 75 bp single-end reads.

#### Bioinformatic Analyzes

Analyzes were carried out on a local Galaxy instance (http://bioinfo.univ-perp.fr). First, the quality of the sequences were checked by FastQC/MultiQC, adapter sequences were removed with Trim Galore! and reads were aligned to version 9 of the *S. mansoni* genome (*schistosoma_mansoni*.*PRJEA36577*.*WBPS17*.*genomic*.*fa*) with permission from https://parasite.wormbase.org/)^27^ using Bowtie2 evoking sensitive end-to-end. Uniquely aligned reads were retained using the Bowtie tag”: “XS:”. PCR duplicates were removed with SamTools RmDup. The number of aligned unique reads was downsampled to 4.7 Mio reads per library using Picard Tools (Suppl. table 2). Differential analysis was done with ChromstaR ^28^with bin size 1000 bp and step size 500 bp.

Annotations came from *schistosoma_mansoni*.*PRJEA36577*.*WBPS17*.*genes*.*gff3*. For metagene analyses, the gene feature was retained and 4898 genes of the forward strand were used.

Differential chromatin states were detected with ChromstaR default parameters.

Peakcalling was performed also with MACS2 and default parameters, followed by MACS2 bdgbroadcall, with 2 and 5 as –cutoff value. Differential HP1 enrichment was detected by DiffBind.^29^ IGV was used for visual inspection.

The same procedures were applied to *D*.*melanogaster* data based on *dmel_r6*.*06_FB2015_03_gtf_dmel-all-r6*.*06*.*gff3* and the corresponding genome fasta file. Details in Suppl. table 2.

#### *In vivo* studies

In vivo study was carried out with 15 female Balb mice divided into 3 groups. Ethical statement at the University of Campinas was under number CEUA protocol (Comissão de Ética no Uso de Animais # 6064-1/2022). Each group contained 5 animals and each animal was infected with 100 3-day-old schistosomula (Groups 1-3). The quantity of schistosomula and cercariae per animal was determined according to the study by Vilar and collaborators^30^. The infection of the groups was subcutaneous, however, in groups 1 and 2, represented by SmHP1 and dsmCherry, respectively, the schistosomula were previously incubated in culture for 3 days with dsRNA. The third group was also kept in culture for 3 days, however, in the absence of dsRNA. For each group, 100 schistosomula were inoculated per animal. Schistosomula was conducted as previously described^31^. Briefly, BH (Belo Horizonte) lineage of the *S. mansoni* was used and schistosomula were transformed by tail break and several RPMI washes for tail removal. After 3 washes, 200 schistosomula were counted and distributed equally in culture plates with 2ml of Medium 169 (Atená, Biotechnology, Campinas, Brazil), described by^32^, supplemented with hormones and fetal bovine serum, 30 μg of dsRNA and kept in CO2 atmosphere at 37°C for 3 days.

All dsRNAs were done using the T7RiboMAX Express RNAi System (Promega, Belo Horizonte, Brazil). Specifically, the oligonucleotides for amplification HP1 gene (T7 + Forward **TAATACGACTCACTATAGGG**TGTGGAAGAGTCAGCTGGT and T7 + reverse **TAATACGACTCACTATAGGG**AGGGTCGATTTTCAGGTGTG), containing T7 promotor for dsRNA synthesis. Gene expressions were evaluated in the QuantStudio qPCR System (Thermo), using the HP1 primers (Primer Forward (CGTCACTCAGTTCAGACAGC) e primer reverse (CTCTTCCACACTCACGGGTA)) and endogenous SmEIF4E (Smp_001500) as described previously^33^, and using the 3 days-schistosomula condition as gene expression calibrator^34^.

Infected animals with wild type and dsRNA were weighted and Kato Katz^20^ were done one week before perfusion. Perfusion was done as described by^35^. For histology, liver tissues were fixed in 10% formaldehyde and fixed in paraffin block and tissue slices were cut and colored by Masson’s trichrome. All granulomas present in 10 random fields of the histological section of each animal were quantified. The images were captured using a photomicroscope using Leica® LAS EZ4 HD software. The total area of the granuloma was measured using ImageJ 1.53t software.

### End Matter

#### Author Contributions and Notes

F.J.C and C.G designed research, C.G. N.S.T. M.V.B. A.R. and T.M.F.M. performed research, C.G analyzed ChIP-seq data, N.S.T. and analyzed *in vivo* results; and F.J.C., C.G and NST wrote the paper.

The authors declare no conflict of interest.

## Acknowledgments

With the support of LabEx CeMEB, an ANR “Investissements d’avenir” program (ANR-10-LABX-04-01) through mobility grants to NST and FJC, and a mobility grant of the Défi RIVOC of the Region Occitanie to FJC. This study is set within the framework of the « Laboratoire d’Excellence (LabEx) » TULIP (ANR-10-LABX-41). Experimental work was done on the Environmental epigenomics platform of the LabEx CeMEB. *D*.*melanogaster* embryos were a generous gift of Bernd Schüttengruber, IGP Montpellier, France. FJC is supported by FAPESP 2021/14982-6 and NST Ph.D. studies is supported by CAPES fellowship supplied by Animal Biology post-graduation program. We thank the Bio-Environment platform (University of Perpignan Via Domitia) and Jean-François Allienne for support in library preparation and sequencing

---

**Supplementary table 1:**
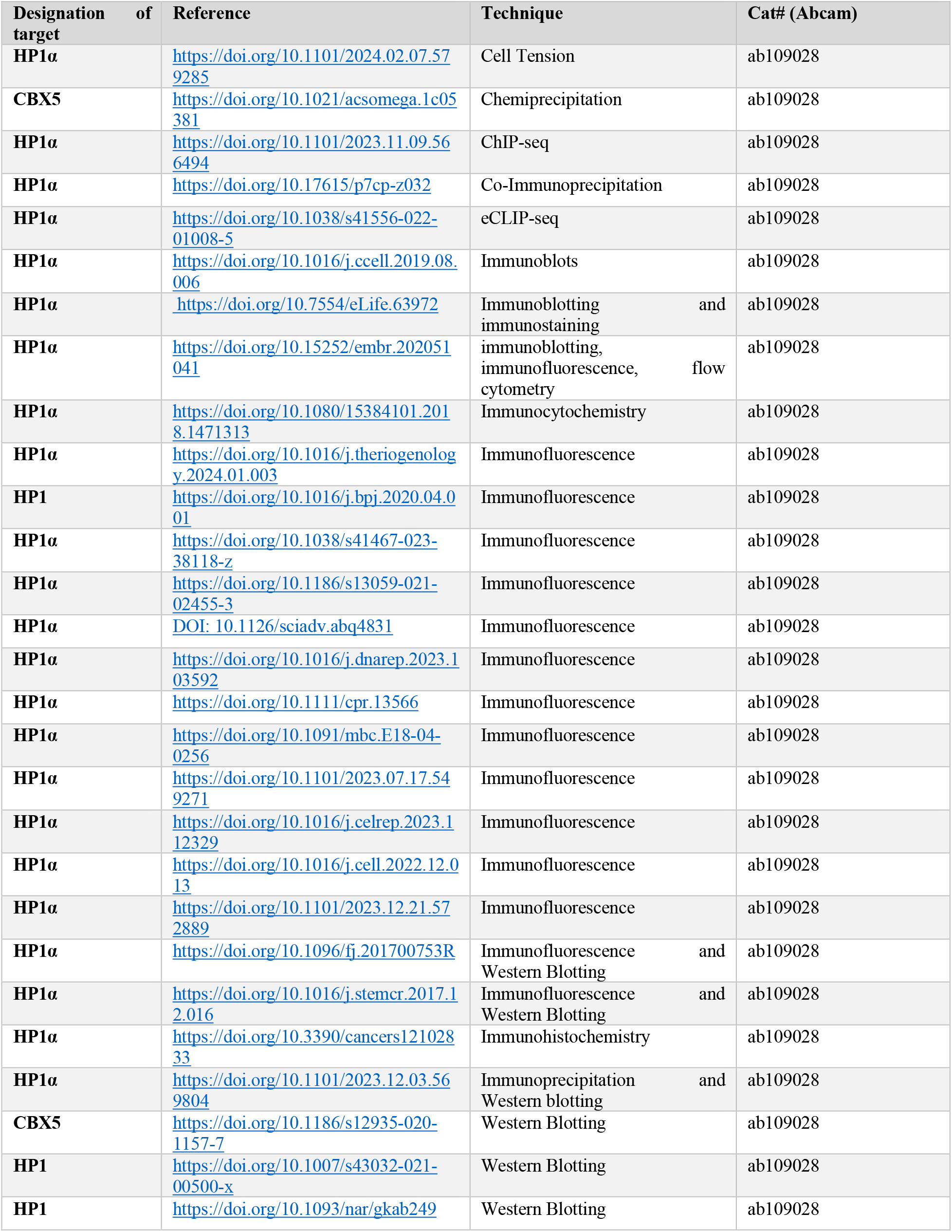
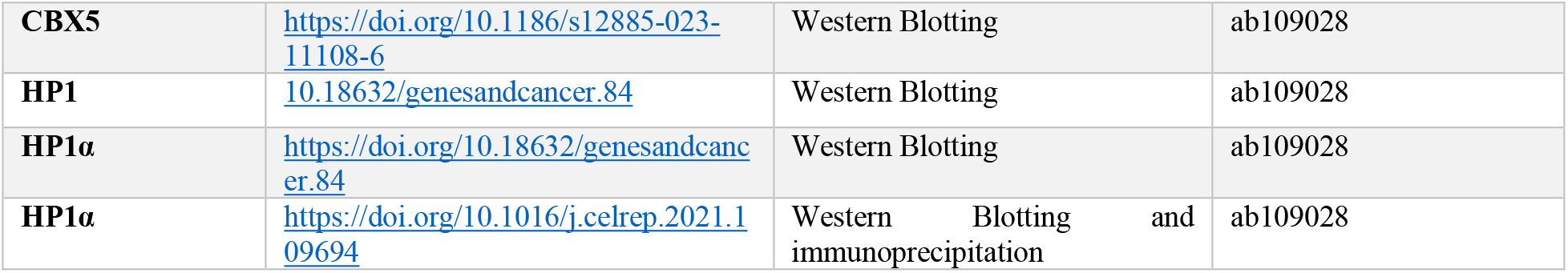
previous use of antiHP1 antbody Abcam ab109028.

**Supplementary table 2:** Detailed sequencing and alignment data.

| Sample | Supplier Name | file name | reads R1 | adapter trimming | Alignment rate | after pick unique and RmDup | in % | Down-sample to |
| --- | --- | --- | --- | --- | --- | --- | --- | --- |
| **Ce 1** | cercaria ChIP HP1 - replica 1 | Ce1_S25_L001_R1_001_fastq.gz | 27,584,702 | 100% | 85.75% | 8,539,009 | 30.96 % | 4,700,000 |
| **Ce 2** | cercaria ChIP HP1 - replica 2 | Ce2_S26_L001_R1_001_fastq.gz | 31,243,784 | 100% | 90.78% | 10,362,799 | 33.17 % | 4,700,000 |
| **Ce 3** | cercaria ChIP HP1 - replica 3 | Ce3_S27_L001_R1_001_fastq.gz | 30,792,782 | 100% | 81.45% | 8,857,714 | 28.77 % | 4,700,000 |
| **Ce-In** | input cercariae undiluted Tn5 | InNa_S31_L001_R1_001_fastq.gz | 14,441,656 | 100% | 96.04% | 9,316,089 | 64.51 % | 4,700,000 |
| **Sp 1** | sporocyst ChIP HP1 - replica 1 | Sp1_S28_L001_R1_001_fastq.gz | 34,896,078 | 100% | 59.46% | 7,340,487 | 21.04 % | 4,700,000 |
| **Sp 2** | sporocyst ChIP HP1 - replica 2 | Sp2_S29_L001_R1_001_fastq.gz | 35,630,630 | 100% | 61.87% | 7,967,706 | 22.36 % | 4,700,000 |
| **Sp 3** | sporocyst ChIP HP1 - replica 3 | Sp3_S30_L001_R1_001_fastq.gz | 29,439,162 | 100% | 44.01% | 4,719,547 | 16.03 % | 4,700,000 |
| **Sp-In** | input tagmentation library sporocysts SmDFO in BgGUA | Sp-In_S7_L001_R1_001_fastq.gz | 188,264,064 | 100% | 57.44% | 16,001,570 | 8.50% | 4,700,000 |
| **Dm2** | D.melanogaster embryos HP1 ChIPmentation rep 2 | Dm2_S9_L001_R1_001_fastq.gz | 157,680,688 | 100% | 83.54% | 18,515,265 | 11.74 % | 16,000,000 |
| **Dm3** | D.melanogaster embryos HP1 ChIPmentation rep 3 | Dm3_S10_L001_R1_001_fastq.gz | 185,909,252 | 100% | 87.05% | 22,123,566 | 11.90 % | 16,000,000 |
| **Dm-In** | D.melanogaster embryos HP1 Input | Dm-In_S11_L001_R1_001_fastq.gz | 173,301,784 | 100% | 95.84% | 20,385,745 | 11.76 % | 16,000,000 |
| **Dm-adult1** | D.melanogaster adults HP1 ChIP rep 1 from SRA | SRR7817540.fastq.gz | 27,849,221 | 100% | 91.26% | 12,676,124 | 45.52 % | 10,000,000 |
| **Dm-adult2** | D.melanogaster adults HP1 ChIP rep 2 from SRA | SRR7817541.fastq.gz | 30,240,617 | 100% | 91.42% | 15,525,830 | 51.34 % | 10,000,000 |
| **Dm-adult3** | D.melanogaster adults HP1 ChIP rep 3 from SRA | SRR7817542.fastq.gz | 37,549,684 | 100% | 89.03% | 10,173,605 | 27.09 % | 10,000,000 |
| **Dm-input** | D.melanogaster adults HP1 Input from SRA | SRR7817573.fastq.gz | 23,583,211 | 100% | 95.44% | 17,566,290 | 74.49% | 10,000,000 |
| **Dm-embryo1** | D.melanogaster embryo HP1 ChIP rep 1 from SRA | SRR11952662.fastq.gz | 19,894,682 | 100% | 94.63% | 12,376,186 | 62.21% | 10,000,000 |
| **Dm-embryo2** | D.melanogaster embryo HP1 ChIP rep 2 from SRA | SRR11952664.fastq.gz | 46,050,252 | 100% | 91.61% | 25,623,510 | 55.64% | 10,000,000 |
| **Dm-embryo-In** | D.melanogaster embryo HP1 Input from SRA | SRR11952661.fastq.gz | 27,346,764 | 100% | 95.32% | 19,767,614 | 72.29% | 10,000,000 |

**Supplementary Figure 1.**
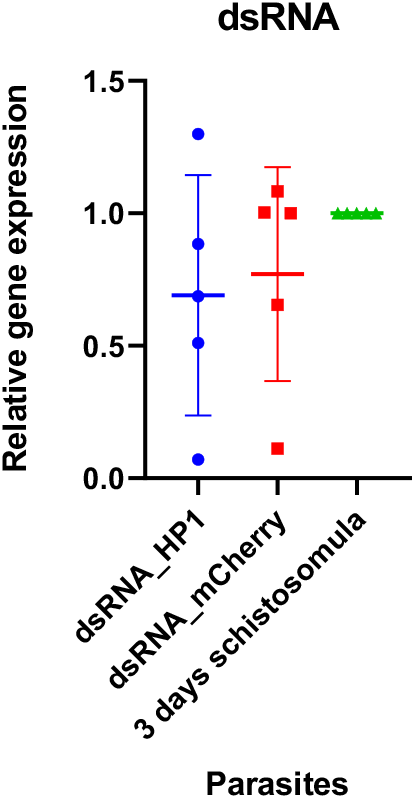
Relative gene expression of dsRNA schistosomula cultures compared to expression of wild type schistosomula. The gene expression was calculated by delta CT method^34^, using wild type schistosomula as gene expression calibrator and the endogenous gene was SmEIF4E (Smp_001500) as described previously^33^

